# Computation of stationary distributions in stochastic models of cellular processes with molecular memory

**DOI:** 10.1101/521575

**Authors:** Jiajun Zhang, Tianshou Zhou

## Abstract

**Abstract:** Modeling stochastic dynamics of intracellular processes has long rested on Markovian (i.e., memoryless) hypothesis. However, many of these processes are non-Markovian (i.e., memorial) due to, e.g., small reaction steps involved in synthesis or degradation of a macroscopic molecule. When interrogating aspects of a cellular network by experimental measurements (e.g., by singlemolecule and single-cell measurement technologies) of network components, a key need is to develop efficient approaches to simulate and compute joint distributions of these components. To cope with this computational challenge, we develop two efficient algorithms: stationary generalized Gillespie algorithm and stationary generalized finite state projection, both being established based on a stationary generalized chemical master equation. We show how these algorithms can be combined in a streamlined procedure for evaluation of non-Markovian effects in a general cellular network. Stationary distributions are evaluated in two models of constitutive and bursty gene expressions as well as a model of genetic toggle switch, each considering molecular memory. Our approach significantly expands the capability of stochastic simulation to investigate gene regulatory network dynamics, which has the potential to advance both understanding of molecular systems biology and design of synthetic circuits.

**Author summary:** Cellular systems are driven by interactions between subsystems via time-stamped discrete events, involving numerous components and reaction steps and spanning several time scales. Such biochemical reactions are subject to inherent noise due to the small numbers of molecules. Also, they could involve several small steps, creating a memory between individual events. Delineating these molecular stochasticity and memory of biomolecular networks are continuing challenges for molecular systems biology. We present a novel approach to compute the probability distribution in stochastic models of cellular processes with molecular memory based on stationary generalized chemical master equation. We map a stochastic system with memory onto a Markovian model with effective reaction propensity functions. This formulation enables us to efficiently develop algorithms under the Markovian framework, and thus systematically analyze how molecular memories regulate stochastic behaviors of biomolecular networks. Here we propose two representative algorithms: stationary generalized Gillespie algorithm and stationary generalized finite state projection algorithm. The former generate realizations with Monte Carlo simulation, but the later compute approximations of the probability distribution by solving a truncated version of stochastic process. Our approach is demonstrated by applying it to three different examples from systems biology: generalized birth-death process, a stochastic toggle switch model, and a 3-stage gene expression model.

## 1. Introduction

Cellular processes are biochemical and hence noisy: in each cell, the numbers or concentrations of species molecules are subject to random fluctuations due to the small numbers of these molecules or environmental perturbations or both. This stochasticity, which has been verified by singlemolecule experiments and can have a significant impact on cellular functioning, has been intensively studied in the past few decades [1,2]. For a biochemical reaction system, if the stochastic motion of the reactants is assumed to be uninfluenced by previous states and only by the current state [3,4], then the stochastic effect can be well revealed by analyzing the chemical master equation (CME) [3-5]. This Markovian assumption has also led to important successes in the description of many stochastic processes on networks [3-8].

However, as a general rule, the dynamics of a given reactive species results from its interactions with the environment and cannot be described as a Markov process. Indeed, although the evolution of the set of all microscopic degrees of freedom of a biochemical system is Markovian, the dynamics restricted to the species only is not [9,10] (Fig.1 (A)). This is typically the case for a promoter cycle, whose non-Markovian dynamics result from the fact that the promoter gradually becomes mature through a series of inactive states (with several hidden Poisson steps) before being activated, leading to narrowly distributed gestation periods between transcription windows [11]. More generally, the expressions of most genes are controlled in a complex manner involving several repressors, transcription factors and mediators, as well as chromatin remodeling or changes in supercoiling. These unspecified regulations or other processes can be modelled by non-exponential waiting time distributions [11-15]. Other experimentally observed examples of non-Markovian dynamics include the human activity patterns [16], natural phenomena [17], biological processes [18], biochemical reactions [19, 20], etc. Examples listed in Table 1 indicate that waiting-time distributions between biochemical events are not limited to exponential distributions but may be diverse. Indeed, the time scales of ultrafast biochemical processes are of the same order as the time scale of the bath they interact with, leading to failure of the Markov approximation.

**Table 1.** Literature survey on waiting-time distributions.

| Description | Timescale | Distribution | References |
| --- | --- | --- | --- |
| Transcription activation time | minute | exponential | 19,20 |
| Transcription inactivation time | hour | phase-type | 19,20 |
| Enzymatic reaction time | millisecond | exponential/phase-type | 21 |
| Lambda induction in bacteria (lysis) | minute~hour | Gamma/phase-type | 22 |
| Transcription initiation time | minute | Gamma/phase-type | 23 |
| Cytokine secretion onset time | hour | Gamma/phase-type | 24 |
| ESC state residence time | hour | Gamma/phase-type | 18 |
| Cell-type specific waiting time | day | Fréchet type | 25 |

**Fig 1.**
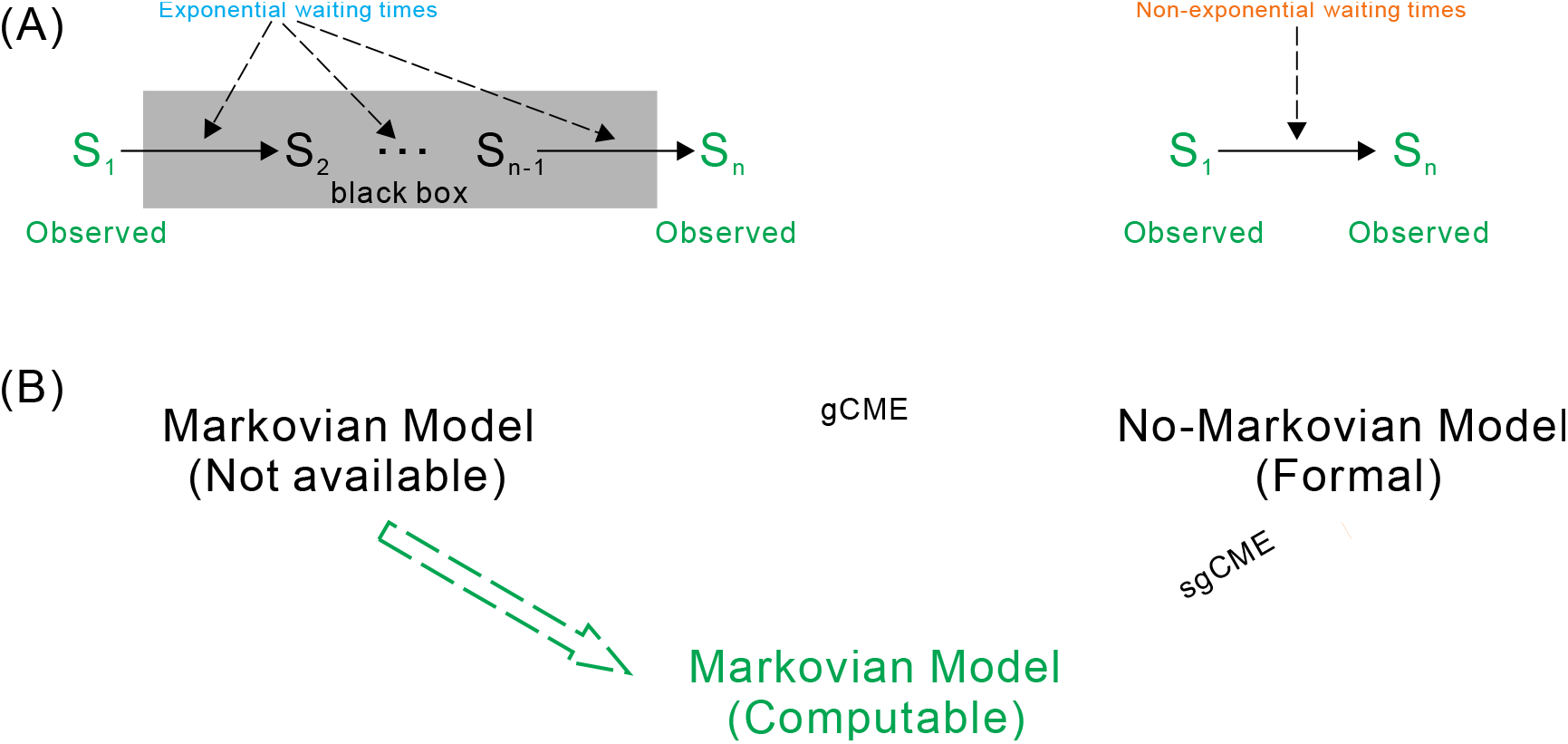
A framework for computing steady-state probability distributions of stochastic reaction processes with molecular memory. (A) Map a multi-step biochemical process onto a single-step reaction process with non-exponential waiting time. (B) Convert a non-Markovian problem into a Markovian problem.

An important step towards a general understanding of non-Markovian dynamics was recently made based on the continuous time random walk (CTRW) theory. Namely, a generalized chemical master equation (gCME) was derived, which can account for non-exponential inter-reaction times and the resulting non-Markovian character of reaction dynamics in time [26,27]. However, this equation is rather formal, and the “memory functions” involved are implicitly expressed by waiting time distributions. This leads to notorious difficulties in analysis and simulation, greatly limiting applications of the gCME. An alternative method is the kinetic Monte Carlo approach, which instead seeks either exact or approximate realizations of the underlying non-Markov process. Earlier, along the line of CTRW, Gillespie developed a stochastic simulation algorithm of random walk with residence time depending on transition probability rates [28], which is actually a prototype of the famous Gillespie algorithm (GA) [29]. Based on the renewal theory, Boguñá and colleagues extended the GA to general renewal processes [30]. Subsequently, Vestergaard and Génois developed a temporal GA, which can simulate non-Markovian dynamics [31]. Recently, Masuda and Rocha proposed a Laplace GA, which can be applied to renewal processes where survival functions are monotonic [32]. In a word, these numerical methods generate realizations enough to obtain statistics for pivotal events. Unfortunately, in many cases, biologically important events occur rarely when reaction waiting times span multiple time scales or/and follow non-exponential distributions, implying that the previous methods are unfeasible in this case. It is an urgent need but is still a significant challenge to develop effective methods to simulate and compute the stochastic dynamics of non-Markovian reaction processes.

As a fact of matter, the emergence of measurement techniques such as single-cell fluorescent microscopy [33], flow and mass cytometry [34], single-cell qPCR [35] and single-cell RNA-seq [36] currently allow for access to increasingly rich data on approximately steady-state distributions of interested molecular species and waiting time distributions of key reaction events. Indeed, steady-state distribution is an important quantity needed to characterize the stationary behavior of many biochemical network systems. To assess how experimental data can be informative, it is also needed to calculate or simulate aspects of steady-state distribution [37-41].

Here we propose two efficient stochastic simulation algorithms: stationary generalized GA (sgGA) that generates realizations with Monte Carlo simulation, and stationary generalized finite state projection (sgFSP) that computes approximate probability distributions. These algorithms are established based on a stationary gCME (sgCME) for a general biomolecular network with arbitrary waiting time distributions. While a key point of the sgCME is that it converts a computationally challenging non-Markovian problem into a computable Markovian one (Fig.1 (B)) by introducing an effective transition rate for every reaction in the network [27], our algorithms are designed to sufficiently exploit this characteristic and hence can be used not only in simulation and analysis of non-Markovian dynamics but also in discovery of biological knowledge underlying non-Markovian effects in complex cellular network systems.

The paper is organized as follows. First, we introduce a general cellular network model with any waiting-time distributions between biochemical reactions. Then, we model this network with a gCME expressed in Laplace transforms and derive a sgCME where effective transition rates decoding non-Markovian effects are explicitly given. Next, we propose two efficient algorithms: sgGA and sgFSP. As an illustration, we have applied these methods to the analysis of a well-characterized birth-death process and investigate the memory effect on stochastic gene expression. Furthermore, we also demonstrated the applicability of these methods to two more complex biochemical network: a stochastic toggle switch model, and a 3-stage stochastic gene expression model. Finally, we revisit derivation of the most common waiting time distributions in reaction systems and give their physical explanations, with the aim to help the reader’s understanding.

## 2. A general theory for reaction networks

Consider a reaction network that models a general cellular process. Assume that this network contains *N* different species (denoted by *X_j_, j* = 1, 2, ⋯, *N*), which participate in *M* different reactions of the form

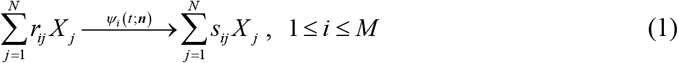

where parameters *r_ij_* and *s_ij_* are stoichiometric coefficients, vector ***n*** = (*n*_1_, ⋯, *n_N_*)^T^ represents the microscopic state of the underlying system with *n_j_* being the molecule number of reactive species *X_j_* and **T** being transpose, and *ψ_i_* (*t*;***n***) be the probability density function (PDF) of the inter-reaction waiting time of they *i*th reaction (see interpretation below). For convenience, let differences *ν_ij_* = *s_ij_* − *r_ij_* be stored in the *M × N* stoichiometric matrix **S**. Note that if the waiting time of all reactions follow geometric distributions, the generalized biochemical reaction network is reduced to the common biochemical reaction network.

Let *P*(*t*;***n***) represent the probability that the system is at state ***n*** at time *t*, and *φ*(*t*;***n***) be the joint PDF of the *i*th reaction happening and the reaction waiting time being *t*, which can be expressed as 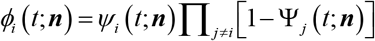. Introduce memory function *M_i_* (*t;**n***) for the *i*th reaction, where the memory is created if inter-reaction waiting time distribution *ψ_i_* (*t*;***n***) is non-exponentail. Following the standard derivation of the Montroll-Weiss equation [42-44] in combination with the idea of chemical continuous time random walks (CTRW) [26,27], we can derive the following gCME in the sense of Laplace transform

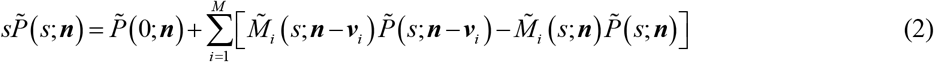

where ***v**_i_* = (*ν*_*i*1_, ⋯, *ν_iN_*)^*T*^, and 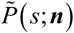 and 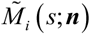 are Lapalace transforms of *P*(*t*;***n***) and *M_i_*(*t*;***n***), respectively.

Interestingly, if we define 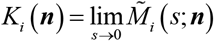 and, then we can show [27]

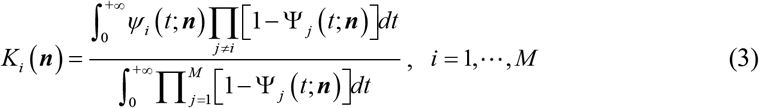

where 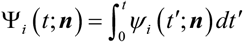 is the cumulative distribution function. Function *K_i_*(***n***) will be called the effective transition rate of the *i*th reaction. Different from reaction propensity function *a_i_*(***n***) in the Markovian case, which is a polymomial of ***n***, function *K_i_*(***n***) may be a rational function of ***n***. Note that effective transition rate (***n***) decodes the effect of non-Markovity in the original reaction system.

Now, we assume that the stationary distribution of the underlying process exists, denoted by *P*(***n***). Then, based on the chemical CRTW theory and effective reaction propensity functions *K_i_*(***n***), we can derive the following sgCME in compact form as [27]

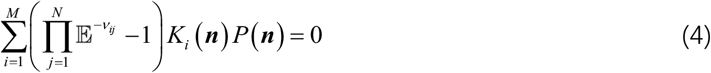

where 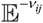 is a step operator with the operation rule below: 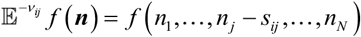 for any function *f*(***n***). If we construct an equivalent reaction network with *K_i_*(***n***) taken as reaction propensity functions, then Eq. (4) has two direct and important implications: (i) the original non-Markovian issue is transformed to an equivalent Markovian one; (ii) although the temporal dynamics of individual reactants would be non-Markovian in the original system, the set of all microscopic degrees of freedom of a reaction system finally evolves to be Markovian, regardless of both the topology of the underlying network and the form of inter-reaction time or/and delay distributions.

## 3. Two efficient algorithms

Based on the above formulation, i.e., Eq. (4) with Eq. (3), we now easily establish numerical algorithms. Here we present two efficient algorithms: sgGA and sgFSP. The former generates realizations with Monte Carlo simulation whereas the later computes an approximate probability distribution. In spite of natural extensions of the corresponding classical algorithms, sgGA and sgFSP are the first algorithms which can efficiently solve a mathematically intractable non-Markovian issue.

### 3.1 Generalized Gillespie algorithm

Recall that traditional GA introduces two random numbers: one for modeling which reaction will happen and the other for modeling when the next reaction will happen. The GA can provide statistically exact methods for simulating stochastic dynamics as interacting sequences of discrete events [29]. However, this algorithm exploits the hypothesis that inter-event times follow exponentially distributions. In the present study, we relax this hypothesis and propose a so-called sgGA based on Eq. (4), which is an extended version of the GA but can be applied to stochastic analysis of a general biochemical network with arbitrary waiting-time distributions. It is worth pointing out that many other extensions of the GA can also be established in cases of non-Markovian reactions.

Similar to the traditional GA, we draw two random numbers *r*_1_ and *r*_2_ from the uniform distribution in the unit interval, and set

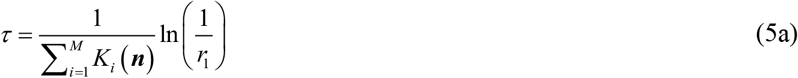

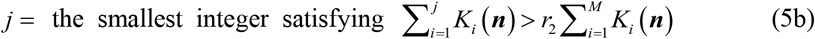

where *K_i_*(***n***) is calculated according to Eq. (3). Then, the sgGA, which constructs an exact numerical realization of the non-Markovian process ***n***(*t*), is stated below

**0**. Initialize the time *t = t*_0_ and the system’s state ***n** = **n***_0_.
**1**. With the system in state ***n*** at time *t*_0_, evaluate all the *K_i_*(***n***) and their sum 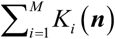.
**2**. Generate values of *τ* and *j* using Eq. 5(a) and 5(b).
**3**. Update the system’s state based on the selected reaction and by replacing *t* ← *t* + *τ* and ***n*** ← ***n*** + (*ν*_1*i*_, *ν*_2*i*_, ⋯, *ν_Ni_*)^T^.
**4**. Record trajectories ***n***(*t*) as desired. Return to Step 1, or else, end simulation.

After discarding the early part of the trajectories (the burn-in phase), the remaining realization values are used to estimate the steady-state probability distribution and related statistics.

### 3.2 Generalized finite state projection algorithm

The FSP method developed initially by Munsky and Khammash has been very successful in solving the CME even for relatively large systems [45]. Based on the idea of FSP, here we propose a sgFSP, which enables to compute stationary probability distributions in systems of non-Markovian biochemical networks.

First, the sgCME (i.e., Eq. 4) can be rewritten in the following form:

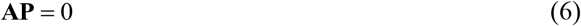

where all possible states space 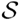 are enumerated as a vector (***n***_1_, ***n***_2_, ⋯)^T^ and **P** is a vector of the stationary probability distributions, i.e., **P**(*t*) = (*P*(*t*;***n***_1_), *P*(*t*;***n***_2_), ⋯)^T^. The *i*th diagonal element of matrix **A** is negative with magnitude equal to the sum of the effective transition rates of reactions that leave the *i*th state, i.e., 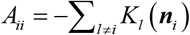, and off-diagonal element *A_ij_* is positive with magnitude being *K_l_*(***n**_j_*) if there is a reaction *l* ∈ {1, ⋯, *L*} such that ***n**_j_* = ***n**_i_* + (*ν*_*i*1_, *ν*_2*i*_, ⋯, *ν_iN_*)^T^ and zero otherwise. In other words, the state reaction matrix **A** comprises of elements:

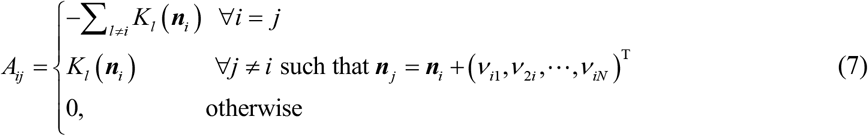

We point out that if the number of the states is infinite or extremely large, the sgCME is very difficult to solve, but one can get a good approximation of the solution by making use of truncation methods such as finite state projection [45], sliding window [46], finite buffer method [47,48] and quantized tensor trains methods [49]. In these methods, truncation is made to guarantee both that the number of states retained is small enough and that the truncated equation is efficiently solved. On contrary, if the number of states retained is very large, the truncated equation is still difficult to solve.

Then, we adopt finite state projection as a foundation to construct a sgFSP. Let 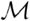 denote a Markov chain on the state set 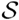, whose master equation is given by *∂*_t_**P**(*t*) = **AP**(*t*). Let 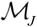 denote a reduced Markov chain, which comprises of the state set 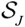 plus a single absorbing state 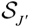 (referring to Fig. 2(B)). The stationary probability distribution satisfying the master equation of 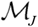 is computed according to

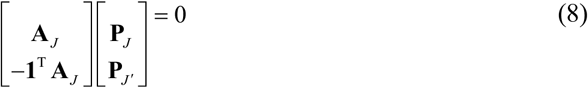

where **1**^T^ = (1, 1, ⋯, 1) is a row vector with the same dimension as that of **A**_*J*_, **P**_*J*_, and **P**_*J′*_, are the stationary probability distributions corresponding to state sets 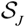 and 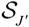, respectively.

**Fig 2.**
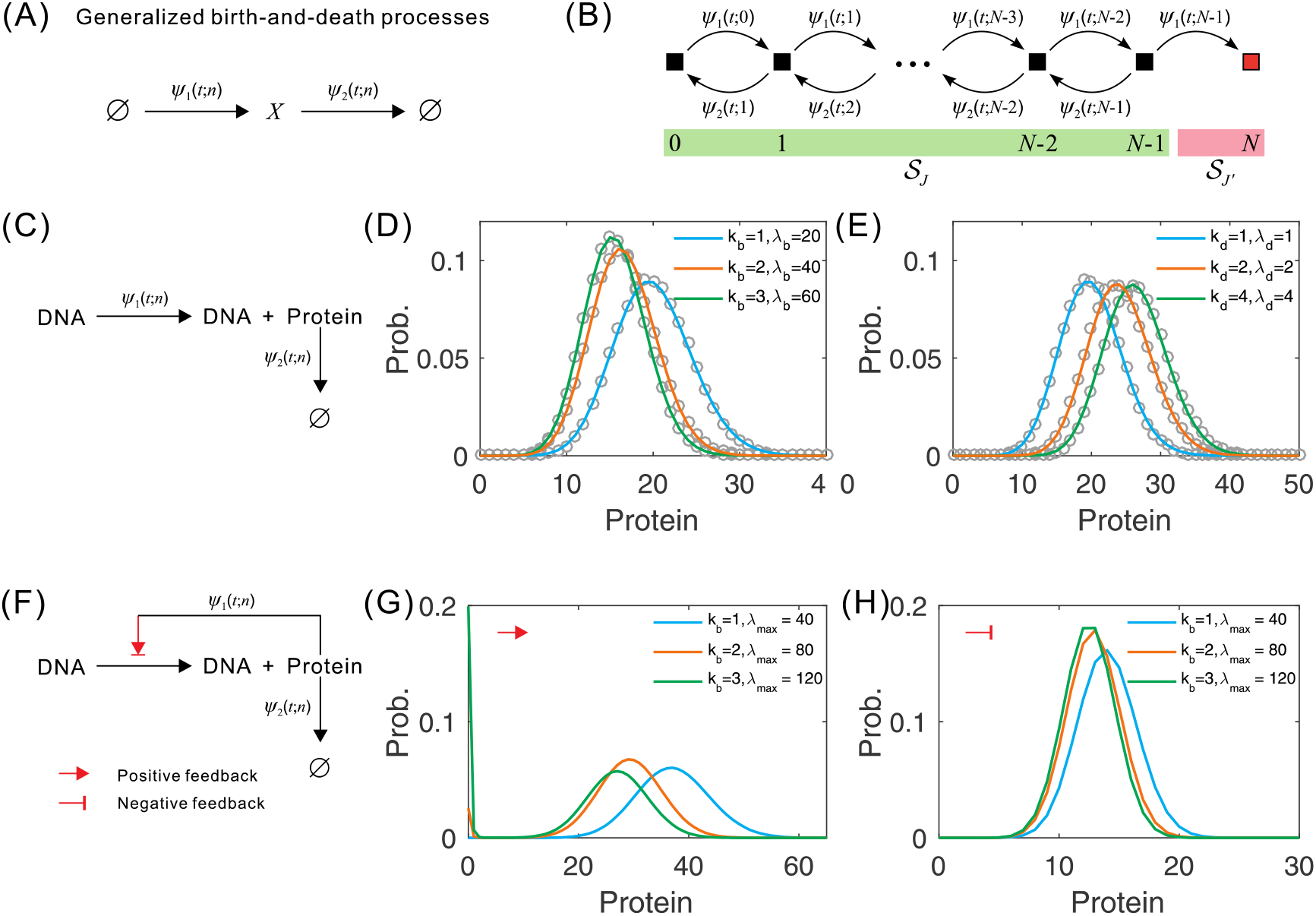
(A) Schematic of generalized birth-death process. (B) A reduced Markov chain with a single absorbing state 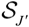. (C) A constitutive gene expression model, where (D) shows protein distributions for different *k_b_*, and (E) show protein distributions for different *k_d_*. (F) A gene self-regulatory model, where (G) shows protein distributions for different *k_b_* and (H) shows protein distributions for different *k_d_*. Some parameter values are set as: (D) *k_d_* = 1, *λ_d_* = 1, (E) *k_b_* = 1, *λ_b_* = 20, (F) *K* = 10, *H* = −2, *ε* = 0.1, *k_d_* = 1, *λ_d_* = 1 and (G) *K* = 10, *H* = 2, *ε* = 0.1, *k_d_* = 1, *λ_d_* = 1.

## 4. Implementations

In order to show the effectiveness of sgGA and sgFSP, here we analyze three representative examples: a generalized birth-death model, a generalized toggle switch model, and an on-off model of gene expression, each considering molecular memory.

### 4.1 A model of birth and death with molecular memory

The first example to be analyzed is a birth-death model, which is a natural and formal framework for modeling a vast variety of biological processes such as population dynamics, speciation, genome evolution [27,50]. Consider a birth-death network consisting of the following two non-Markovian reactions (Fig. 2(A))

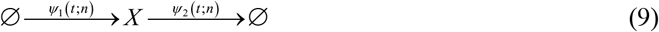

where two waiting time distributions are set as 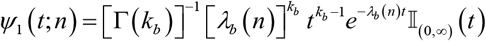 and 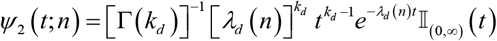. Symbol 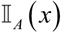 is an indicator function of set *A*, i.e., 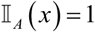 if *x* ∈ *A* and 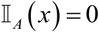 otherwise. Symbol Γ(·) is the common Gamma function. In numerical simulation, we limit ours consideration to the case of Gamma distributions, although our approach may be applied to other kinds of distributions suggested in the literature [51] (additional explanations can be found in the next section).

It is worth pointing out that the above formulation contains the stochastic model of a gene’s constitutive expression as its special case, where protein synthesis corresponds to birth while protein degradation to death [52,53]. Here, we consider the following two cases of gene expression (referring to Figure 2(C) and (F)):

**Case 1** Gene expression without feedback

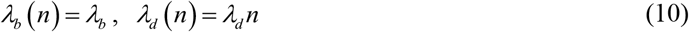

**Case 2** Gene expression with feedback

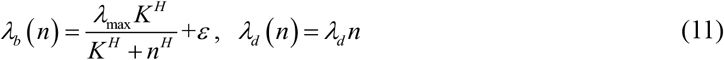

where *λ*_max_ is the maximum transcription rate, *K* is the equilibrium constant of the transcription factor binding to its DNA binding site, and *ε* represents a small leakiness of the promoter. Hill coefficient *H* > 0 corresponds to negative feedback and *H* < 0 to positive feedback.

In order to show how sgFSP is performed, we set 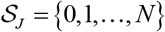 (Fig. 2(B)). In addition, denote **P** = (*P*(0), ⋯, *P*(*N*))^T^, where the conservative condition 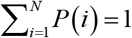 must hold. Introduce a matrix of the form

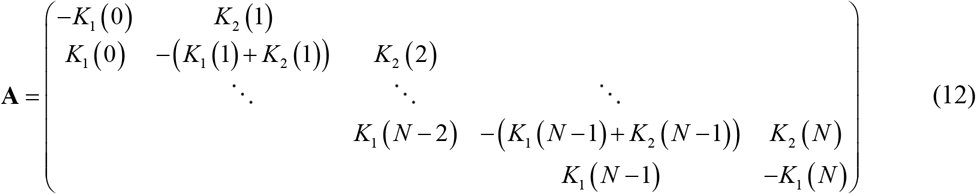

By solving the algebraic equation **AP** = 0 with 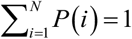, we can obtain a numerical **P**.

For clarity, we distinguish the following four cases to demonstrate numerical results: (1) *ψ*_1_(*t*;*n*) is a Gamma distribution whereas *ψ*_2_(*t*;*n*) is an exponential distribution (i.e., *k_d_* = 1) in **case 1** (Fig. 2(D)); (2) *ψ*_2_(*t*;*n*) is a Gamma distribution whereas *ψ*_1_(*t*;*n*) is an exponential distribution (i.e., *k_b_* = 1) in **case 1** (Fig. 2(E)); (3) *ψ*_1_(*t*;***n***) is a Gamma distribution for positive feedback regulation whereas *ψ*_1_(*t*;*n*) is an exponential distribution in **case 2** (Fig. 2(G)); (4) *ψ*_1_(*t*;*n*) is a Gamma distribution for negative feedback regulation whereas *ψ*_2_(*t*;*n*) is an exponential distribution in **case 2** (Fig. 2(H)). In all the cases, we set *k_b_*/*λ_b_*, *k_d_*/*λ_d_* and *k_b_*/*λ*_max_ as constants, so that the average waiting times remain unchanged. Recall that a Gamma distribution can model a multistep process. Therefore, integer parameter *k_b_* (or *k_d_*) represents the step number in our case.

Numerical results are demonstrated in Fig. 2. In Fig. 2(D)(E), lines correspond to distributions obtained by sgFSP and empty circles to distributions obtained by sgGA. From these two panels, we observe that the two obtained kinds of distributions match remarkably well, implying that the two methods are equivalent. Therefore, we focus on the performance of sgFSP in the following. We also observe that protein distributions shifts toward the direction that *k_b_* increases (Fig. 2(D)(G)(H)), but toward the opposite direction (Fig.2 (E)). Interestingly, protein populations demonstrate bimodal distribution when *k_b_* > 1 (Fig.2 (E)), implying that molecular memory can induce population diversity.

### 4.2 Gene model of toggle switch with molecular memory

Recall that a toggle switch network (Fig. 3(A)) can model the cross-repression between the determinants of different cellular states, which can result in a definite choice between two outcomes [54-56]. Conventional models of genetic toggle switch consider exponential waiting time distributions. However, the expression of a gene in general involves a multi-steps process. Indeed, transcriptional repressor monomer (A or B) binds first to dimers and then to specific DNA sequences near the promoter, repressing the production of transcriptional repressor monomer (B or A). This multi-step process can result in non-exponential waiting times and hence create molecular memories between reaction events. Here, we consider a stochastic model of genetic toggle switch with memories, which consists of the following four reactions (or referring to Fig. 3(B)):

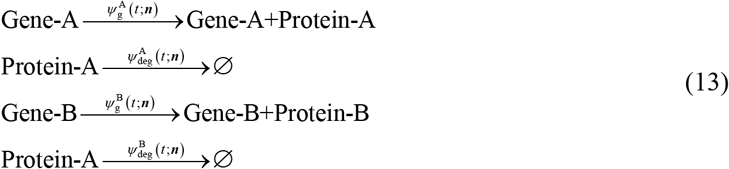

where 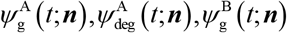 and 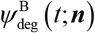 are waiting time distributions. We assume that these distributions are:

1. Transcription process for gene A: 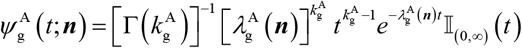 with 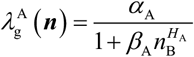;
2. Degradation process for gene A: 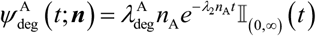;
3. Transcription process for gene B: 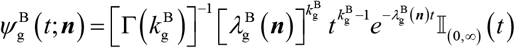 with 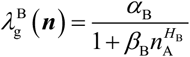;
4. Degradation process for gene B: 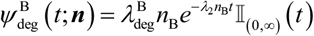.

**Fig 3.**
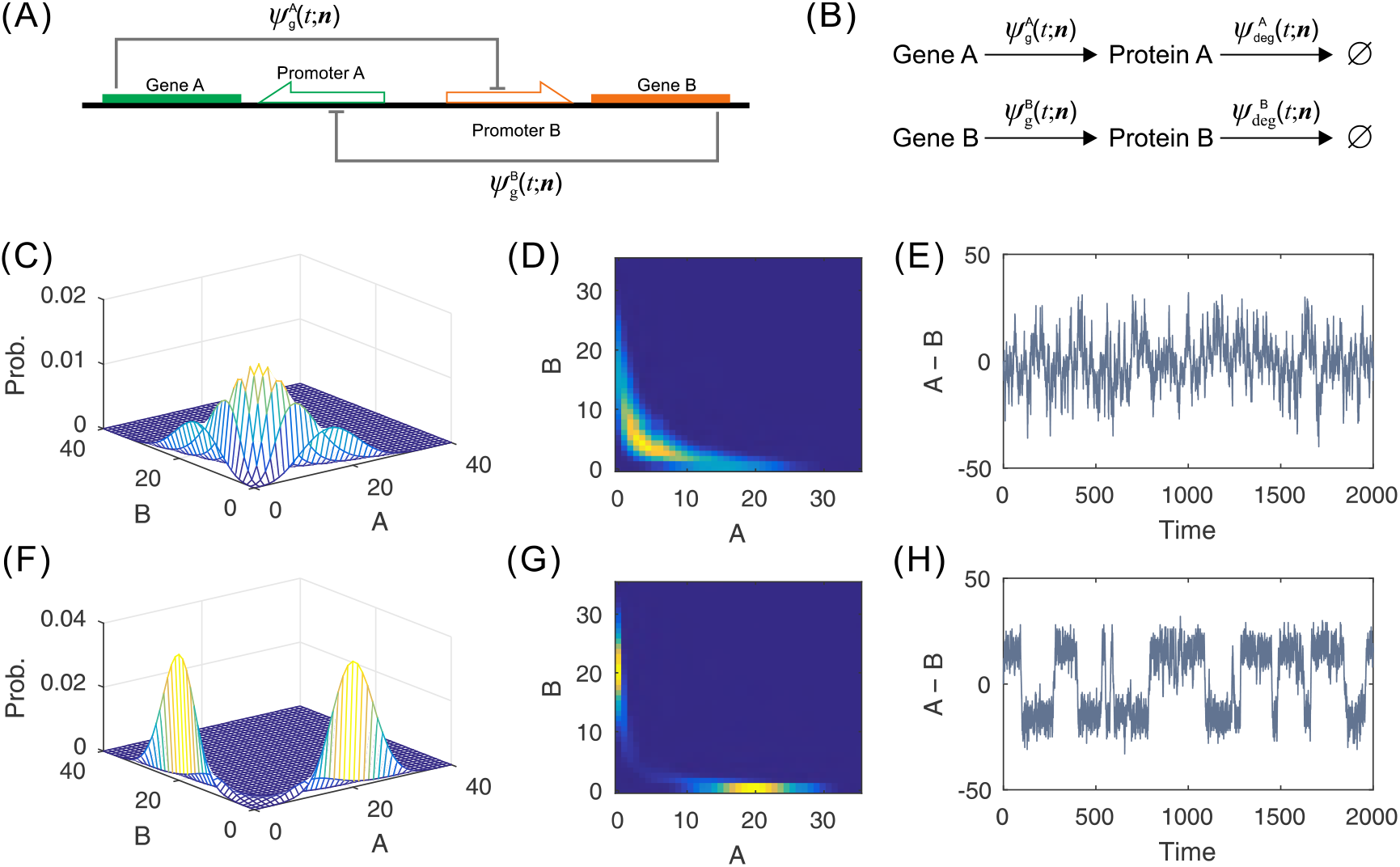
(A) Schematic for a model of toggle switch with memory, where two genes are repressed to each other. (B) Four reactions corresponding to (A), where waiting times for synthesis and degradation of each protein follow distributions, respectively. Default parameter values are taken as: 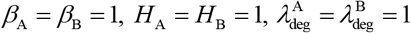. (C) and (F) Joint distributions of proteins A and B, obtained by sgFSP; (D) and (G) Heat maps in the plane of protein A and B; (E) and (H) Time series of the difference between the levels of protein A and B, obtained by sgGA. (C)-(E) correspond to exponential waiting times, where parameter values are set as 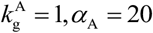; (F)-(H) correspond to non-exponential waiting times, where parameter values are set as 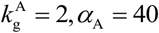.

Note that 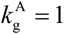 and 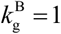 1 correspond to the Markovian case whereas 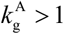 or 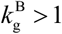 corresponds to the non-Markovian case. In numerical simulations, we always set 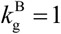.

Numerical results are demonstrated in Fig. 3, where Fig. 3(C)-(E) correspond to the case of exponential waiting times whereas Fig. 3(F)-(H) to the case of non-exponential waiting times. We observe that if the waiting time for synthesis of protein A follows an exponential distribution (i.e., if 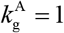 is set), the steady-state joint distribution of protein A and B is unimodal, referring to Fig. 3(C)(D). However, if the waiting time for synthesis of protein A follows an non-exponential distribution (i.e., if 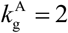 is set), the steady-state joint distribution of protein A and B is bimodal, referring to Fig. 3(F)(G). To examine the time dependence of the populations of two proteins in a single cell, we first perform stochastic simulations with sgGA and then calculate the difference between the levels of proteins A and B. The numerical results are shown in Fig. 3(E)(H). Comparing Fig. 3(E) with Fig. 3 (H), we find that two switching states occur only in the case of non-exponential waiting times or memory. In a word, we conclude from Fig. 3 that molecular memory can induce bimodal distributions in the toggle switch model described in Fig. 3(A) or Fig. 3(B).

### 4.3 On-off model of gene expression with molecular memory

The 3-stage model of gene expression [12-15,57,58] can find its prototypes in prokaryotic and eukaryotic cells. This model, which can capture many essential features of transcriptional biology, has extensively been used and studied. The basic hypothesis of the model is that the gene promoter exists either in an activated state that produces mRNA or in a repressed state that is unproductive, and that an mRNA undergoes a competition between translation and degradation. In traditional studies, it is also assumed that every chemical reaction involved happens probabilistically at a fixed rate, i.e., waiting times between reaction events are assumed to follow exponential distributions. However, biochemical events do not necessarily occur in single steps of individual molecules. For example, a promoter’s gradual maturing through a series of inactive states (many of which have been unspecified) before activation can result in non-exponential distributed gestation periods between transcription windows. Also for example, mRNAs or proteins’ senescing through a series of states (many of which have also been unspecified) before they are finally degraded can also lead to non-exponential waiting times.

In order to model various possible regulations involved in gene expression, we replace exponential waiting time distributions with non-exponential waiting time distributions for transitions between promoter states, transcription, translation and degradation. Thus, the traditional 3-stage model of gene expression is modified as (also referring to Fig. 4(A)):

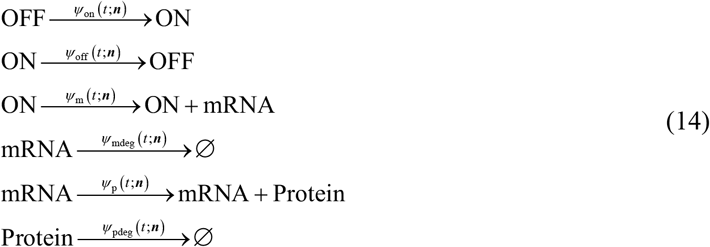

where OFF denotes an inactive promoter state, ON denotes an active promoter state, and the first two reactions describe the promoter switch between its OFF and ON states. The ON-state DNA can generate mRNAs that are further translated into proteins and are eventually degraded. Functions *ψ*_on_ (*t*;***n***), *ψ*_off_ (*t*;***n***), *ψ*_m_ (*t*;***n***), *ψ*_mdeg_(*t*;***n***), *ψ*_p_ (*t*; ***n***) and *ψ*_pdeg_ (*t*;***n***) in Eq. (14), which are waiting time distributions, can describe complex regulatory processes. For convenience, we denote by ***n*** = (*on, off, m, p*)^T^ the state vector of the gene system, where *on, off, m, p* represent the numbers of DNA molecules at off and on states, the numbers of mRNA and protein molecules, respectively. We assume that all these waiting time distributions are Gamma distributions, that is,

1. Promoter switch process: 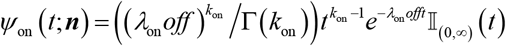, 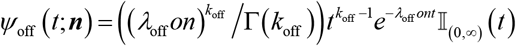.
2. Transcription process: 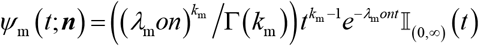.
3. mRNA degradation process: 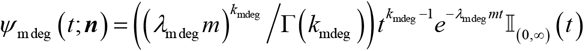.
4. Translation process: 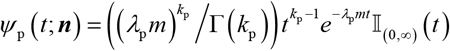.
5. Protein degradation process: 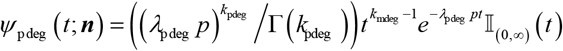.

**Fig 4.**
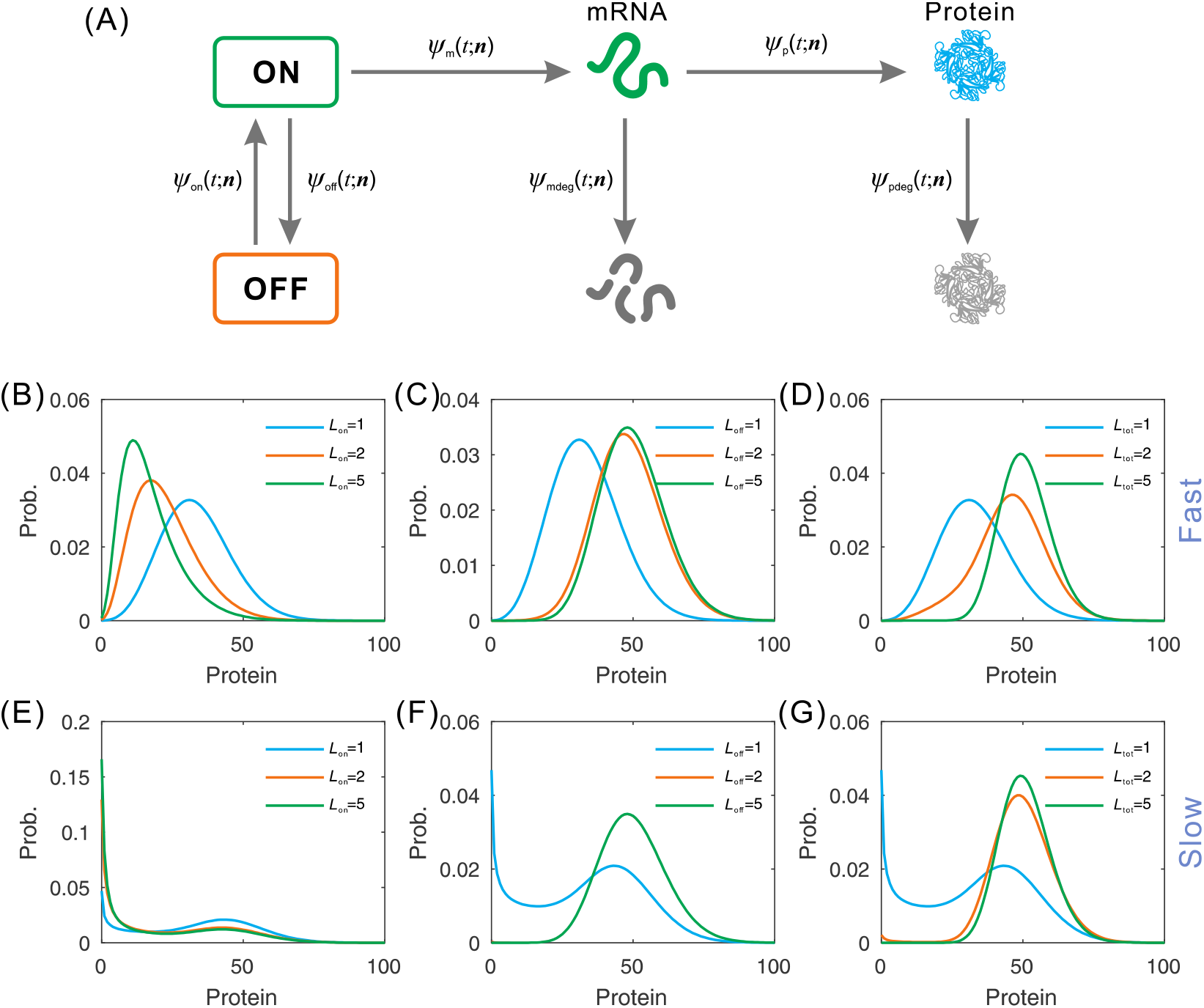
(A) Schematic for a 3-stage model of gene expression with molecular memory. (B)-(G) Protein distributions obtained by sgFSP, where default parameter values are set as following *k*_on_ = 1, *λ*_on_ = 1 (corresponding to fast switch) or *λ*_on_ = 0.1 (corresponding to slow switch), *k*_off_ = 1, *λ*_off_ = *λ*_off_ = 0.5 (fast switch) or *λ*_off_ = 0.05 (slow switch), *k*_m_ = 1, *λ*_m_ = 5, *k*_mdeg_ = 1, *λ*_mdeg_ = 1, *k*_p_ = 1, *λ*_p_ = 2, *k*_pdeg_ = 1, *λ*_pdeg_ = 0.2. In (B) and (E), *k*_on_ = *L*_on_, *λ*_on_ = *L*_on_ (fast switch) and *λ*_on_ = 0.1*L*_on_ (slow switch). In (C) and (F), *k*_off_ = *L*_off_, *λ*_off_ = 0.5*L*_off_ (fast switch) and *λ*_off_ = 0.05*L*_off_ (slow switch). In (D) and (G), all the default parameter values are multiplied by *L*_tot_.

As is well known, the relative rate of the constant gene-state transition rates has significant impact on gene expression dynamics. Here, we compute protein distributions in two dynamic regimes: ‘fast’ gene-state dynamics (possibly due to fast binding and unbinding of transcription factors)) and ‘slow’ gene-state dynamics (possibly due to slower changes in chromatin configuration). Note that parameters *L*_on_ and *L*_off_ in the model represent the step numbers of multistep processes, which result in Gamma distributed waiting times. In simulation, the average waiting times is kept constant. Numerical results are shown in Fig.4 (B)-(G). From these panels, we observe that whichever “fast” and “slow” switches, protein distributions shifts to smaller number density as *L*_on_ is increased (Fig.4 (B)(E)), but shifts higher number density as *L*_off_ (*L*_tot_) is increased (Fig.4 (C)(D)(F)(G))). Interestingly, protein populations tend to limit distributions as *L*_off_ (*L*_off_, *L*_tot_) is increased.

## 5. Discussion

A key challenge in stochastic modeling of cellular networks is in raising computational efficiency involving the jointing effects of stochastic fluctuations (or the noise) and molecular memory. We have developed a novel methodological approach, which is based on sgCME, to compute steady-state probability distributions in systems of general cellular reaction processes with molecular memory. While the sgCME would open a new way to address this classically intractable problem from a Markovian viewpoint, the approach can develop a set of efficient simulation methods. Here, we have proposed a Monte Carlo simulation algorithm (i.e., sgGA) to generate realizations of stochastic trajectories, and an approximate algorithm (i.e., sgFSP) to directly compute steady-state probability distributions. Analysis of three representative exemplars has verified the effectiveness of these algorithms.

Several potential weaknesses yet computational challenges with our methods exist both in simulation of transient stochastic dynamics and in choice of an appropriate waiting time distribution. First, our methods rest on sgCME. Although this equation can reveal long time behavior of stochastic process with memory, but lacks moderately transient dynamics. To overcome this difficulty, there are two possible roadmaps: One is that one may take the whole line back to the gCME formulation, numerical integrate the corresponding integro-differential equation, and store the whole historical data; And another is to resort to Monte Carlo simulation. Future extensions of the algorithms are under way. Second, the waiting time distribution selection is essentially a statistic inference problem in systems biology. Large amount of experimental data imply that Gamma distribution and phase-type distribution are good choices. The specifics are outside the scope of this article and will be discussed elsewhere.

We stress that our framework opens a door to analyze and simulate stochastic behaviors of the systems of cellular processes with molecular memory. The presented formulation enables us to efficiently develop algorithms under a Markovian framework with effective transition rates instead of reaction propensity functions. And our theory is general in that it can deal with biochemical networks of arbitrary topology and arbitrary types of waiting time distributions. Therefore, theoretically, successful analysis methods and simulation algorithms developed in the Markovian framework can apply to non-Markovian problems [59]. In addition, it is worth pointing out that our formalism is not restricted to biochemical networks considered here and can be easily extended to any non-Markovian networks with a finite/infinite set of discrete states, such as epidemic spreading on complex networks [60-63].

Finally, the emergence of measurement techniques currently allow for access to increasingly rich data on approximately steady-state distributions of interested molecular species and waiting time distributions of key reaction events. The presented methods are computationally efficient and scalable. This will facilitate the quantitative mechanistic modelling of complex cellular processes and the exploitation of cell-to-cell variability and molecular memory.

## Appendix: Biophysical foundation of waiting-time distributions

The timing of cellular processes cannot be described often by simple Poisson processes, which are traditionally characterized by exponential distributed waiting time distributions and hence can be modelled by classical chemical master equations. However, many single-peaked and even multimodal waiting-time distributions have been observed empirically, reflecting complexity of cellular processes modelled by biochemical reactions networks. Indeed, non-exponential waiting-time distributions are signatures of relevant intracellular processes without a priori knowledge. A main advantage of gCME capturing effects of non-exponential waiting times is that it can focus on a small number of key measurable events. Here, we derive the most common waiting time distributions of biochemical reactions: exponential distribution, Gamma distribution and phase-type distribution, focusing on their biophysical explanations.

### Exponential distribution

Exponential distributions can characterize the time between events in a Poisson point process. Exponential waiting times between reactions are a pillar of the classical chemical master equation. Although exponential waiting time distributions can be derived in different ways, an approximate analysis of molecular collision times provides a clear physical explanation.

Assume that all molecules are spheres with diameter *D*. If the center of one molecule gets the center of another molecule within a distance *D*, a collision occurs. Specifically, in the next small time interval *dt*, a molecule will sweep out a volume equal to *πD^2^vdt*, in the sense that if another molecule center falls within this volume, a collision will occur in the time interval (*t, t+dt*). *N* with *N*>>1 molecules sweep out this volume for *N* times. For a given molecule center, the probability that it falls within the total volume swept out by the other molecule is then approximately *NπD^2^vdt/V*, where *V* is the total volume and v is the average molecule speed. The probability of no collision for a given molecule is [29],

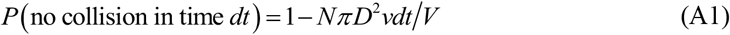

The probability of no collision after *N* time intervals *dt* is

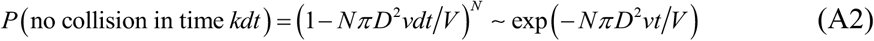

where *t = Ndt*. The probability of a collision is 1 − *P* (no collision in time *Ndt*), and the probability density of *t* can be written

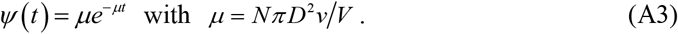

It is the exponential distribution which can be used to modeling the waiting times between single elementary reaction steps. Note that this derivation is based on Markovian assumption and the exponential distribution can also be derived from a master equation [3,4].

### Gamma distribution

In contrast to the times until the next molecular event in single-step processes (elementary reactions), the waiting times between reactions are, in general, not exponentially distributed. If a reaction process goes through a series of (identical) exponential steps, the waiting times will be Gamma distributed [51]. Mathematically, this process can be regarded as a sum of *n* single-step processes, and therefore the over-all waiting time is the *n*-fold convolution

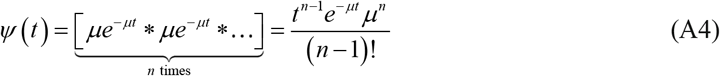

where, * denotes convolution and *ψ*(*t*) is known as the gamma distribution with shape parameter *n* and rate parameter *μ*. If *n* is a positive integer, then the distribution is an Erlang distribution.

### Phase-type distribution

A phase-type distribution is a probability distribution constructed by a convolution or mixture of exponential distributions. It results from a system of one or more inter-related Poisson processes occurring in sequence or phases, such as reversible chemical reactions. For example, models describing enzyme dynamics can be mapped onto phase type models. Phase-type distributions can provide a tractable theoretical framework to understand complex waiting time distributions observed for a variety of different biochemical processes. Formally, the PDF of a phase-type distribution is [51]

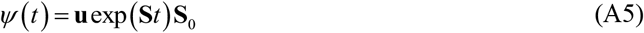

where **S** is an *m × m* matrix, **S**_0_ is a *m* × 1 vector which consisting of the negative sums of the columns of matrix **S**, and **u** = (1, 1, ⋯, 1) is a 1 × *m* vector.

There are a lot of probability distributions involving waiting times, such as Weibull distribution, Pareto distribution, and Mittag-Leffler distribution. We do not tend to enumerate all here.

## Acknowledgments

This work was supported by grants 91530320, 11775314, 11475273, and 11631005 from Natural Science Foundation of P. R. China; 2014CB964703 from Science and Technology Department, P. R. China; 201707010117 from the Science and Technology Program of Guangzhou, P. R. China.

## Author Contributions

Conceived and designed the experiments: JJ, TS. Performed the experiments: JJ. Analyzed the data: JJ. Contributed reagents/materials/analysis tools: JJ, TS. Wrote the paper: JJ, TS.

